# Collateral development and arteriogenesis in hindlimbs of domestic swine after ligation of arterial inflow

**DOI:** 10.1101/491928

**Authors:** Y Gao, NS Patel, S Aravind, M Fuglestad, JS Ungar, CJ Mietus, S Li, GP Casale, II Pipinos, MA Carlson

## Abstract

**Introduction:** The development of collateral vasculature is a key mechanism compensating for arterial occlusions in patients with peripheral artery disease (PAD). We aimed to examine the development of collateral pathways after ligation of native vessels in a porcine model of PAD.

**Methods:** Right hindlimb Ischemia was induced in domestic swine (N=11, male) using two different versions of arterial ligation. Version 1 (N=6) consisted of ligation/division of the right external iliac, profunda femoral and superficial femoral arteries. Version 2 (N=5) consisted of the ligation of Version 1 with additional ligation/division of the right internal iliac artery. Development of collateral pathways was evaluated with standard angiography at baseline (prior to arterial ligation) and at termination (4 weeks later). Relative luminal diameter of the arteries supplying the ischemic right hindlimb were determined by 2D angiography.

**Results:** The dominant collateral pathway that developed after Version 1 ligation connected the right internal iliac artery to the right profunda femoral and then to the right superficial femoral/popliteal artery. Mean luminal diameter of the right internal iliac artery at termination increased by 38% compared to baseline. Two co-dominant collateral pathways developed in Version 2 ligation: (i) from the left profunda femoral artery to the reconstituted right profunda femoral artery; and (ii) from the common internal iliac trunk and the left internal iliac artery to the reconstituted right internal iliac artery, which then supplied the right profunda femoral and then the right superficial femoral/popliteal artery. Mean diameter of the left profunda and the left internal iliac artery increased at termination by 26% and 21%, respectively (p < 0.05).

**Conclusion:** Two versions of hindlimb ischemia induction (right ilio-femoral artery ligation with and without right internal iliac artery ligation) in swine produced differing collateral pathways, along with changes to the diameter of the inflow vessels (i.e., arteriogenesis). Radiographic and anatomical data of the collateral formation in this porcine model has value in investigation of the pathophysiology of hindlimb ischemia, and assessment of angiogenic therapies as potential treatments for PAD.

## INTRODUCTION

Peripheral artery disease (PAD) has a prevalence of 3-10% in the general population, with an impact on functional status, quality of life, and life expectancy [*1–3*]. The vast majority of symptomatic PAD patients present with exercise-associated leg discomfort and ambulatory disability known as intermittent claudication [*1–3*]; only 1-2% of patients present with critical ischemia, including severe pain at rest and/or tissue loss/gangrene. Treatment options for symptomatic PAD include open and percutaneous revascularization but, after referral, only 13% of patients with claudication and 50% of patients with critical limb ischemia undergo a revascularization procedure [*4,5*]. These decreased intervention rates are driven by the presence of complex occlusive disease and/or medical comorbidities which can limit walking ability and/or increase operative risk. For patients not undergoing revascularization, development of collateral blood flow is an important compensatory mechanism to improve blood flow to the ischemic limb [*5,6*], which makes the study of blood vessel growth in response to ischemia an important field of investigation.

The groups of Drs. Brewster and Lefer in collaboration with our group [*7,8*], have demonstrated that two swine strains (with and without comorbidities) can recapitulate the pathophysiology of human PAD, including (i) persistently decreased hindlimb hemodynamics/perfusion, (ii) decreased treadmill performance, (iii) and ischemic myopathy (end-organ damage). The ability of swine to model human PAD prompted us to use a porcine model of PAD to determine: 1) the pathways and size of the collaterals developing over thirty days after two different versions of arterial ligation in the right hindlimb; and 2) the effect of these ligations and resulting collateral development on limb hemodynamics, oxygenation, and muscle histology. The rationale for studying this collateralization phenomenon was to develop a clinically-relevant large-animal platform in which we could test the efficacy of cell-based therapies (or other interventions) in improving perfusion of the ischemic limb.

## MATERIALS AND METHODS

### Animal Welfare Statement

The animals utilized to generate the data for this report were maintained and treated in accordance with the *Guide for the Care and Use of Laboratory Animals* (8^th^ ed.) from the National Research Council and the National Institutes of Health [*9*] and also in accordance with the Animal Welfare Act of the United States (U.S. Code 7, Sections 2131 – 2159). The animal protocol pertaining to this manuscript was approved by the Institutional Animal Care and Use Committee (IACUC) of the VA Nebraska-Western Iowa Health Care System (ID number 00950) and by the IACUC of the University of Nebraska Medical Center (ID number 15-068-07-ET). All procedures were performed in animal facilities approved by the Association for Assessment and Accreditation of Laboratory Animal Care International (AAALAC; www.aaalac.org) and by the Office of Laboratory Animal Welfare of the Public Health Service (grants.nih.gov/grants/olaw/olaw.htm). All surgical procedures were performed under isoflurane anesthesia, and all efforts were made to minimize suffering. Euthanasia was performed in accordance with the AVMA Guidelines [*10*].

### Experimental Subjects

Wild type domestic swine (N = 13; castrated males; age 11-13 wk) were purchased from the Animal Research and Development Center at the University of Nebraska Lincoln (ardc.unl.edu). Swine were housed individually in pens, with multiple subjects per room, and fed ad lib with standard hog feed (Purina Nature’s Match Sow and Pig Complete Feed).

### Experimental Design

Refer to Fig. tf01. The design included an ischemia-induction procedure (i.e., arterial ligation) on day 0, with endpoint measurement and necropsy 30 d later. Two methods of arterial ligation were compared in two nonrandomized groups of domestic swine; the groups were matched for age, size, and sex.

**Fig. tf01.**
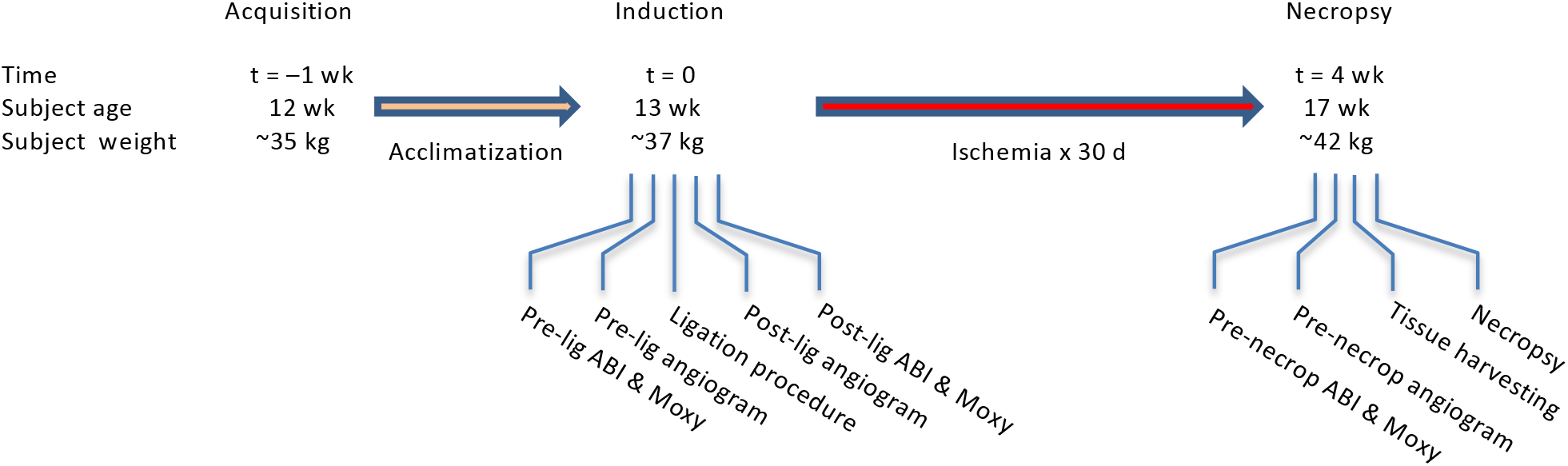
Experimental flow diagram. Domestic swine were 3 mo old at time of acquisition (t = −1 wk).

### Set-Up and Anesthesia[*11*]

Swine were fasted for 24 h prior to the procedure, with free access to water and up to two 500 cc bottles of regular Gatorade^TM^. On day zero, each subject underwent induction with ketamine (Zoetis, 2.2 mg/kg), Telazol^®^ (Zoetis; 4.4 mg/kg) and xylazine (Zoetis; 2.2 mg/kg), given as a single IM injection. Each subject then was weighed and endotracheally intubated. EKG, pulse oximetry, rectal temperature, and lingual end-tidal CO_2_ monitors were placed. The subject rested on a water-circulated warming blanket that was set at 102°F. An auricular intravenous (IV) line was placed. Anesthesia was maintained with isoflurane (0.5-1%) and supplemental oxygen (3-5 L/min) using a Matrx^®^ ventilator (midmark.com). The ventilator rate initially was set at 12-15 breaths per minute with a tidal volume of 10 mL/kg, and subsequently adjusted to maintain the EtCO2 at 40-50 mm Hg. Cotton blankets were placed over non-surgical areas to minimize any decrease in body temperature. Vital signs were continuously recorded to a laptop computer via a Bionet BM5 monitor (www.bionetus.com). A single dose of cefovecin sodium (Zoetis; 8 mg/kg IM) was given before incision.

### Muscle Oximetry & Arterial Indices

Extremity muscle oximetry measurements were measured transcutaneously in the supine position under general anesthesia using a Near Infrared Spectroscopy probe (Moxy device, moxymonitor.com) with the Peripedal software (peripedal.com); see Fig. tf11S. The Moxy device has a noninvasive probe which measures muscle hemoglobin/myoglobin oxygen saturation (StO_2_). In the forelimb, the Moxy probe was placed over the medial biceps brachii of the forelimb (i.e., medial aspect of the flexor mass of the forelimb), proximal to the elbow joint; in the hindlimb, the probe was placed over the medial gastrocnemius muscle. The Moxy index was defined as the ratio of the hindlimb saturation over the higher of the two forelimb saturations. For determination of arterial indices, a pediatric-size sphygmomanometer cuff was applied to each extremity, about 10 cm above the hoof (see Fig. tf11S), and systolic pressure was measured with the aid of a Doppler device (Model 811-B, Parks Medical Doppler; parksmed.com). The hindlimb-forelimb arterial pressure index (“ABI”, meant to be the porcine analog of the clinical ankle-brachial index) was defined as the ratio of hindlimb systolic blood pressure over the higher of the two forelimb systolic pressures.

### Laparotomy, Vessel Exposure, and Angiography

With the subject under general anesthesia and in the supine position, the chest, abdomen, groins, and bilateral lower extremities were shaved with an electric clipper, washed with soap and water, and then prepped using ChloraPrep™ applicators (chlorhexidine gluconate/isopropyl alcohol; BD Medical). Vessel exposure was obtained using a retroperitoneal approach through a right paramedian incision that started to the right and just inferior to the urethral meatus and medial to the right nipple line, and then extended inferiorly across the right inguinal ligament (not divided) and onto the right inner thigh (Fig. tf10S), The abdominal wall layers were incised carefully down to the peritoneum, and then the dissection was performed laterally between the peritoneum and abdominal wall to develop the retroperitoneal space. The peritoneal sac, containing the intraabdominal organs, was retracted superiorly and to the left, which exposed the distal aorta and right-sided pelvic vasculature. In all subjects an arterial line (20-gauge Angiocatheter) was placed in the infrarenal aorta, 4-5 cm above the aortic trifurcation, through a site controlled with a 6-0 polypropylene U-stitch (see Fig. tf05, and the postmortem dissection in Fig. tf09S). This line was used for arterial blood pressure monitoring and angiography. A baseline aortogram with runoff was obtained with injection of 10 mL of Visipaque™ 320 (lodixanol; GE Healthcare) and C-arm fluoroscopy (GE OEC 9900 Elite).

### Arterial Interruption

Refer to Figs. tf05, tf02, and tf09S. Once baseline angiography was obtained, the region of the aortic trifurcation was dissected. In the pig, the distal aorta trifurcates into the right and left external iliac arteries (REIA and LEIA) and the common internal iliac trunk (CIIT); the latter then bifurcates into the right and left internal iliac arteries (RIIA and LIIA). The REIA was dissected down to the bifurcation of the right superficial femoral artery (RSFA) and profunda femoris artery (RPFA). The REIA, right lateral circumflex iliac and lateral circumflex femoral artery (RLCIA and RLCFA, respectively) were suture ligated proximally with 3-0 silk. Distal dissection exposed the bifurcation of the RSFA into the right popliteal and saphena arteries (RPop and RSA). The RSFA was suture ligated distal to the takeoff of the RLCFA. A continuous segment of the REIA-RSFA was then excised (Figs. tf05 and tf02). This series of steps constituted the Version 1 arterial interruption procedure. Version 2 consisted of Version 1 plus ligation of the RIIA. To summarize:

**Fig. tf02.**
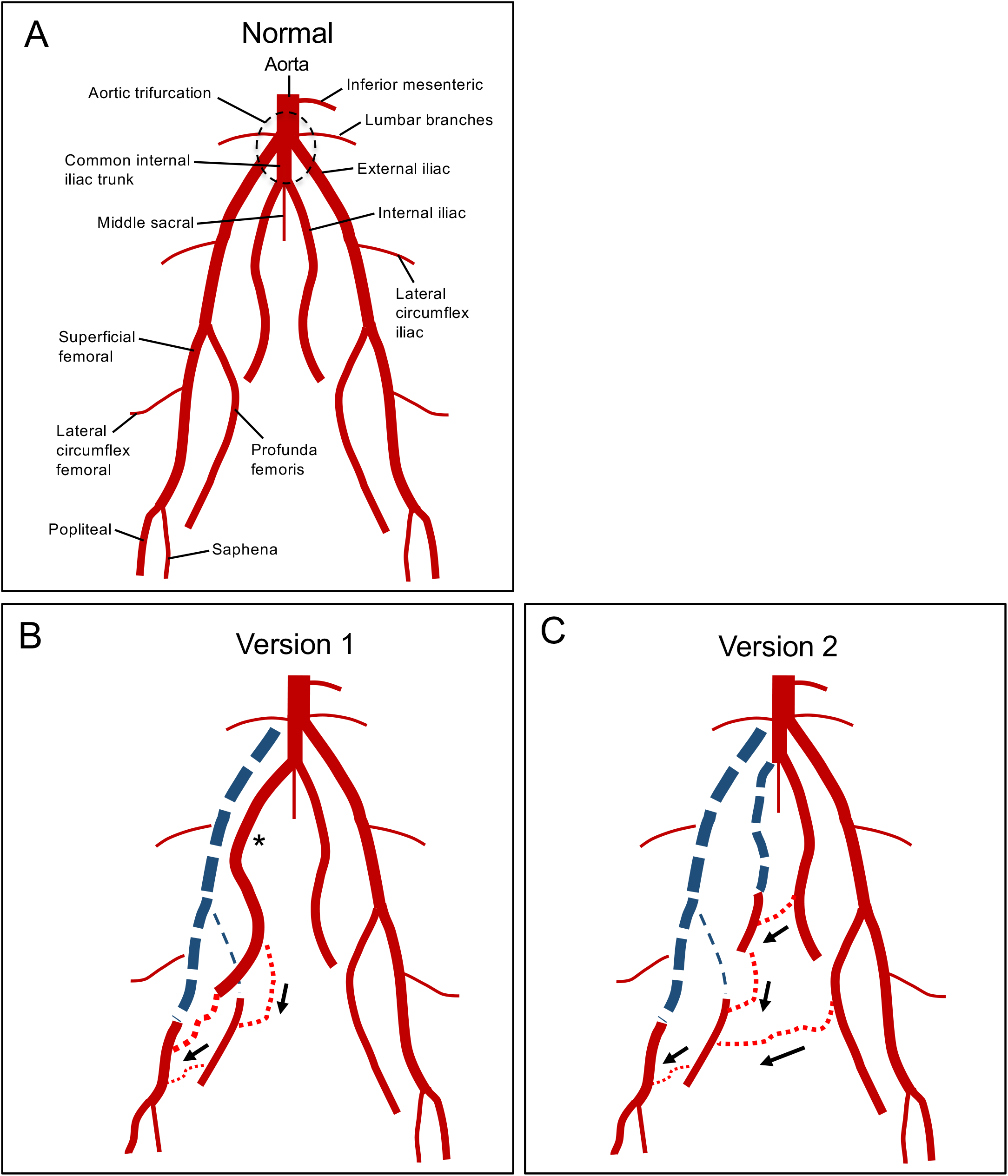
Arterial anatomy of the porcine hindlimb. Normal anatomy. **(B)** Ligation Version 1. The right iliofemoral segment has been ligated and excised (thick dashed blue line), along with the right profunda femoris (thin dashed blue line). One month after ligation, the right internal iliac artery has enlarged (*), and there is collateral flow from this vessel which reconstitutes both the distal superficial femoral artery and the distal right profunda femoris (black arrows). **(C)** Ligation Version 2. In addition to the ligations performed for Version 1, the right internal iliac vessel was ligated. One month after ligation, collaterals from the left profunda femoris reconstitute the right profunda femoris while collaterals from the left internal iliac artery reconstitute the right internal iliac, which then helps reconstitute the right profunda femoris and distal right superficial femoral artery through further collateralization (black arrows).

> Version 1 Interruption (Fig tf02B): ligation of the REIA, RSFA, RPFA, RLCIA, RLCFA and excision of a continuous segment of the REIA-RSFA.
>
> Version 2 Interruption (Fig. tf02C): Version 1 + RIIA ligation

A completion angiogram was performed after arterial interruption was accomplished. The aortic puncture site utilized for angiographic access was then closed by removing the angiocatheter and tying the previously-placed U-stitch. The abdominal incision was closed anatomically in layers, with 2-0 polyglactin 910 in the peritoneum and posterior layers, 0-polydioxanone in the anterior aponeurosis, 3-0 polyglactin 910 in the panniculus carnosus, and 4-0 polyglactin 910 in the skin. Cyanoacrylate glue was applied over the skin incision; no other incisional dressing was applied. The animal’s recovery from anesthesia was monitored until the subject was awake and mobile. Subjects were given half feeds on post-ligation day 1, and were placed back on ad lib feeds on day 2.

### Terminal procedure

Subjects underwent their terminal procedure 4 weeks after arterial interruption. After induction of general endotracheal anesthesia, the right carotid artery was exposed with a cervical cut-down. Hindlimb perfusion measurements (muscle oximetry and arterial indices) were performed as described above. A catheter was then placed into the right common carotid artery and advanced into the infrarenal aorta using the Seldinger technique. An abdominal aortogram with distal runoff was performed. Bilateral hindlimb dissection was performed prior to euthanasia. Biopsies of the medial heads of the bilateral gastrocnemius were harvested prior to euthanasia (Fig. tf13S) and processed as described under “Bright-Field Microscopy”.

### Arterial Diameter Analysis

Each angiogram obtained during the procedure was saved into the Digital Imaging and Communications in Medicine (DICOM^®^) format within the GE OEC 9900 Elite C-arm instrument. Images subsequently were accessed and analyzed using the RadiANT DICOM Viewer (Version 4.0.2; www.radiantviewer.com). The cross-sectional diameter of the distal aorta was defined as 10 units. Relative diameter of each distal artery was then expressed in units of distal aortic diameter.

### Bright-Field Microscopy

Porcine muscle specimens were fixed immediately in cold methacarn fixative for 48 h, transferred into cold 50% ethanol, and then embedded in paraffin. Sections (4 µm) from these blocks were deparaffinized in xylene, rehydrated in water, and then H&E stained. Bright field images were captured with a Leica DMRXA2 microscope configured with a Leica DFC color camera (North Central Instruments; www.ncimicro.com). Sections from archived paraffin blocks of human gastrocnemius muscle from deidentified patients (a PAD patient and a control subject) were similarly processed and imaged. These human specimens were obtained through an IRB protocol utilized in a prior study [*12*].

### Statistical analysis

Continuous data were analyzed using ANOVA within Microsoft Excel; the level of significance was defined as p < 0.05.

## RESULTS

### Perioperative Events

Mean starting weight in subjects who completed the study was 32.8 ± SD 4.0 kg (range = 26.6 – 39.0 kg, N = 6) and 37.0 ± 10.5 kg (range = 35.0 – 55.6 kg, N = 5) for Versions 1 and 2, respectively (p = 0.19, unpaired t-test). A total of 13 subjects underwent the ligation procedure. One pig developed non-reducible rectal prolapse four days after operation and was euthanized; another pig expired on postoperative day one secondary to intestinal ischemia. Necropsy of the latter subject revealed a stomach distended with feed and generalized severe ischemia (purplish discoloration) of the small and large intestine, but otherwise was inconclusive; specifically, all mesenteric vessels were patent. Of the eleven subjects which survived to the scheduled termination (N = 6 for Version 1 and N = 5 for Version 2), two subjects (one of each version) developed clinically-apparent ischemia of the right hoof, manifested by ulcer formation on the weight-bearing region. Their mobility was affected, but each animal could still ambulate and both survived until scheduled termination.

### Arterial Pressure Indices at Rest

One subject’s ABI and Moxy data (Version 1) were missing, so ten subjects underwent analysis of ABI and Moxy data. The ABI was calculated at rest (i.e., under anesthesia) in both hindlimbs on day 0 (pre-ligation and immediately post-ligation), and then again on day 30 (pre-necropsy); see Table 1. In both the ischemic (right) and non-ischemic (left) hindlimb, the mean ABI was not different between the two ligation versions at any individual time point, including at day 30 (unpaired t-tests in Table 1). Immediately post-ligation in the ischemic hindlimb, the ABI dropped to zero for both versions. At 30 d (pre-necropsy), the mean ABI in the ischemic hindlimb had recovered in both ligation versions to ~0.6. In the non-ischemic hindlimb, the ABI did not change significantly over time in either ligation version.

**Table 1.** Hindlimb arterial pressure indices (“ABI”) in Version 1 vs. 2.

**Right (ischemic) hindlimb**
| Version (N) | Day 0 |  | ABI, Day 30 | *ANOVA |
| --- | --- | --- | --- | --- |
|  | ABI, pre | ABI, post |  |  |
| 1 (N = 5) | 1.00 ± 0.22 | 0 | 0.62 ± 0.25 | 0.0001 |
| 2 (N = 5) | 1.24 ± 0.30 | 0 | 0.62 ± 0.35 | 0.0004 |
| **Unpaired t-test | 0.196 | 1.000 | 0.968 |  |

**Left (non-ischemic) hindlimb**
| Version (N) | Day 0 |  | ABI, Day 30 | *ANOVA |
| --- | --- | --- | --- | --- |
|  | ABI, pre | ABI, post |  |  |
| 1 (N = 5) | 1.18 ± 0.23 | 1.02 ± 0.09 | 1.15 ± 0.20 | 0.365 |
| 2 (N = 5) | 1.18 ± 0.30 | 1.09 ± 0.26 | 1.22 ± 0.24 | 0.722 |
| **Unpaired t-test | 0.974 | 0.589 | 0.610 |  |

The reference limb pressure (the “brachial” pressure or denominator in the ABI ratio) was the higher of the two forelimb pressures. ABI was expressed as mean ± standard deviation. Terms “pre” and “post” refer to before and after arterial ligation. The Day 30 ABI was determined immediately prior to necropsy. \*The ANOVA compares ABI of one hindlimb in one version over three time points. \*\*The unpaired t-test compares ABI between two versions at a single time point.

### Muscle hemoglobin/myoglobin oxygen saturation (StO_2_) at Rest

Similar to ABI, the Moxy index (i.e., ratio of hindlimb over forelimb StO_2_), was calculated at rest (i.e., under anesthesia) in both hindlimbs on day 0 (pre-ligation and immediately post-ligation), and then again on day 30 (pre-necropsy); see Table 2. Bar plots of absolute hindlimb StO_2_ values are shown in Supplemental Fig. tf14S. In both the ischemic (right) and non-ischemic (left) hindlimb, the mean Moxy index was not different between the two ligation versions at any individual time point (unpaired t-tests in Table 2), with one exception. At day 30 in the non-ischemic hindlimb, the mean Moxy index was slightly but significantly higher in the Version 2 subjects. Immediately post-ligation in the ischemic hindlimb, the mean Moxy index dropped to ~0.4 for both versions. At 30 d (pre-necropsy), the mean Moxy index in the ischemic hindlimb had recovered in both ligation versions to 0.6 – 0.7. In the non-ischemic hindlimb, the mean Moxy index did not change significantly over time in either ligation version, though there appeared to be a post-ligation trend to a decrease for both ligation versions. Interestingly, the bar plots of the raw Moxy measurements (Fig. tf14S) demonstrated a definite dip in mean StO_2_ immediately post-ligation in the non-ischemic limb for both ligation versions; this decrease was still significant at day 30 for Version 1. Otherwise, the bar plots of the raw Moxy measurements yielded results comparable with the Moxy index data.

**Table 2.**
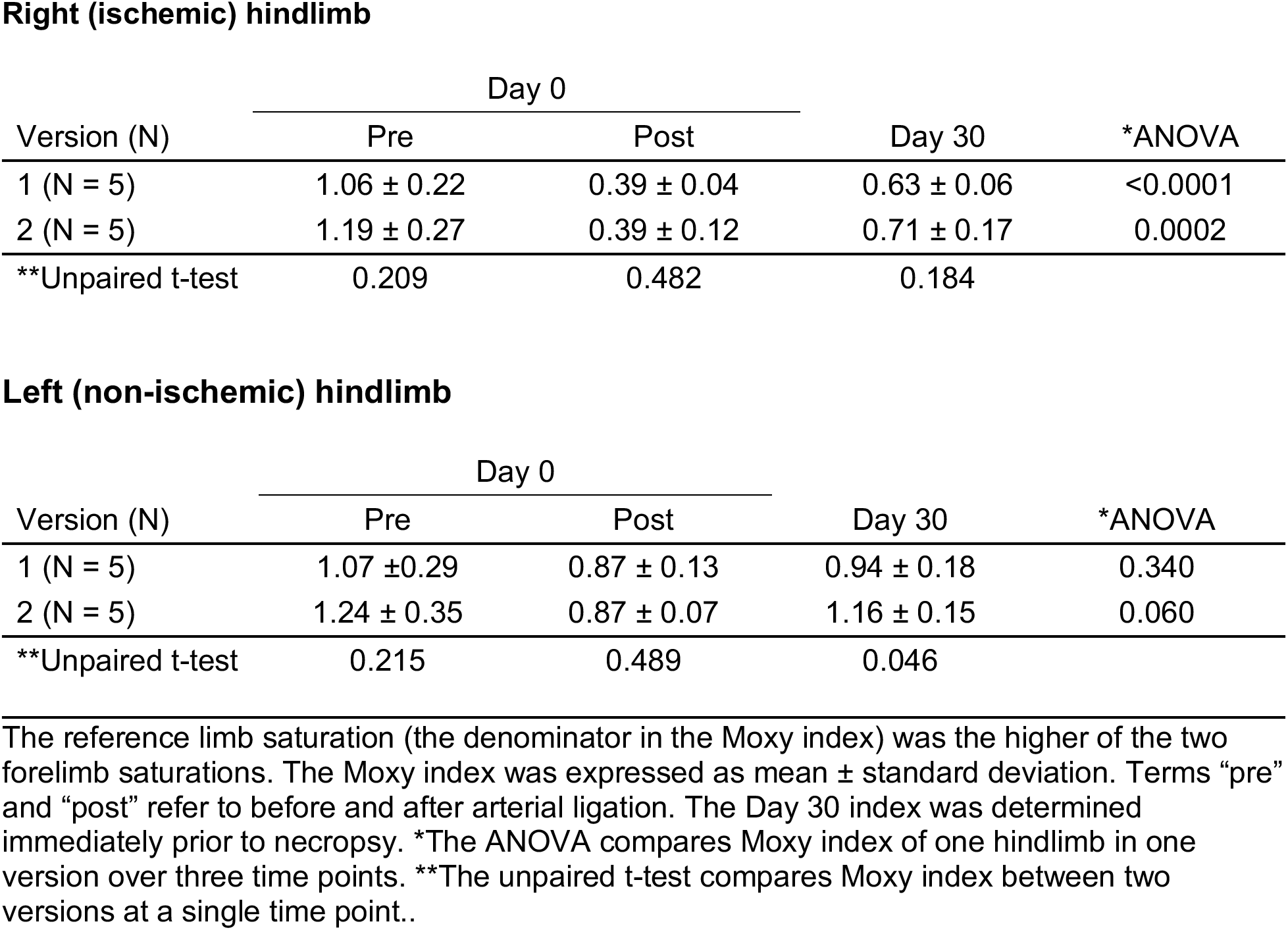
Muscle hemoglobin/myoglobin oxygen saturation (Moxy) indices in Version 1 vs. 2.

**Right (ischemic) hindlimb**
| Version (N) | Day 0 |  | Day 30 | *ANOVA |
| --- | --- | --- | --- | --- |
|  | Pre | Post |  |  |
| 1 (N = 5) | 1.06 ± 0.22 | 0.39 ± 0.04 | 0.63 ± 0.06 | <0.0001 |
| 2 (N = 5) | 1.19 ± 0.27 | 0.39 ± 0.12 | 0.71 ± 0.17 | 0.0002 |
| **Unpaired t-test | 0.209 | 0.482 | 0.184 |  |

**Left (non-ischemic) hindlimb**
| Version (N) | Day 0 |  | Day 30 | *ANOVA |
| --- | --- | --- | --- | --- |
|  | Pre | Post |  |  |
| 1 (N = 5) | 1.07 ± 0.29 | 0.87 ± 0.13 | 0.94 ± 0.18 | 0.340 |
| 2 (N = 5) | 1.24 ± 0.35 | 0.87 ± 0.07 | 1.16 ± 0.15 | 0.060 |
| **Unpaired t-test | 0.215 | 0.489 | 0.046 |  |

The reference limb saturation (the denominator in the Moxy index) was the higher of the two forelimb saturations. The Moxy index was expressed as mean ± standard deviation. Terms “pre” and “post” refer to before and after arterial ligation. The Day 30 index was determined immediately prior to necropsy. \*The ANOVA compares Moxy index of one hindlimb in one version over three time points. \*\*The unpaired t-test compares Moxy index between two versions at a single time point..

### Collateral Pathway Development

Refer to Videos 1-12 in the Supplemental Material. Sample images (video stills) of the pre-ligation (baseline) aortography with runoff for Versions 1 and 2 are shown in Figs. tf04A and tf06A, respectively (see accompanying Videos 1-2 and 7-8, respectively). Immediately after ligation in Version 1, the REIA-RSFA segment was no longer present; there was delayed filling of the RFPA and remaining distal arteries from the nonligated RIIA, with some minor contribution to the distal hindlimb through collaterals involving the RCIA (Fig. tf04B and Videos 3-4). After 30 days of ischemia in Version 1, pre-necropsy angiography (Fig. tf04C) demonstrated that the RIIA was noticeably enlarged. Collateral vessels from the RIIA reconstituted the RPFA, which was followed by RSFA/RPop reconstitution (Video 5-6). The RSFA/RPop was reconstituted either from the RIIA through collaterals to the RLCFA, or through collaterals from the RPFA system. Immediately after ligation in Version 2, both the REIA-RSFA segment and the RIIA were no longer visible (Fig. tf06B; Video 9-10). There was delayed filling of the RIIA and RPFA, but no reconstitution of RSFA or Rpop. After 30 days of ischemia in Version 2, pre-necropsy angiography (Fig. tf06C; Video 11-12) demonstrated the presence of two co-dominant collateral pathways supplying the ischemic right hindlimb: (i) from the LPFA to the reconstituted RPFA; and (ii) from the CIIT and LIIA to the reconstituted RIIA, subsequently supplying the reconstituted RPFA and RSFA/RPop.

**Fig. tf04.**
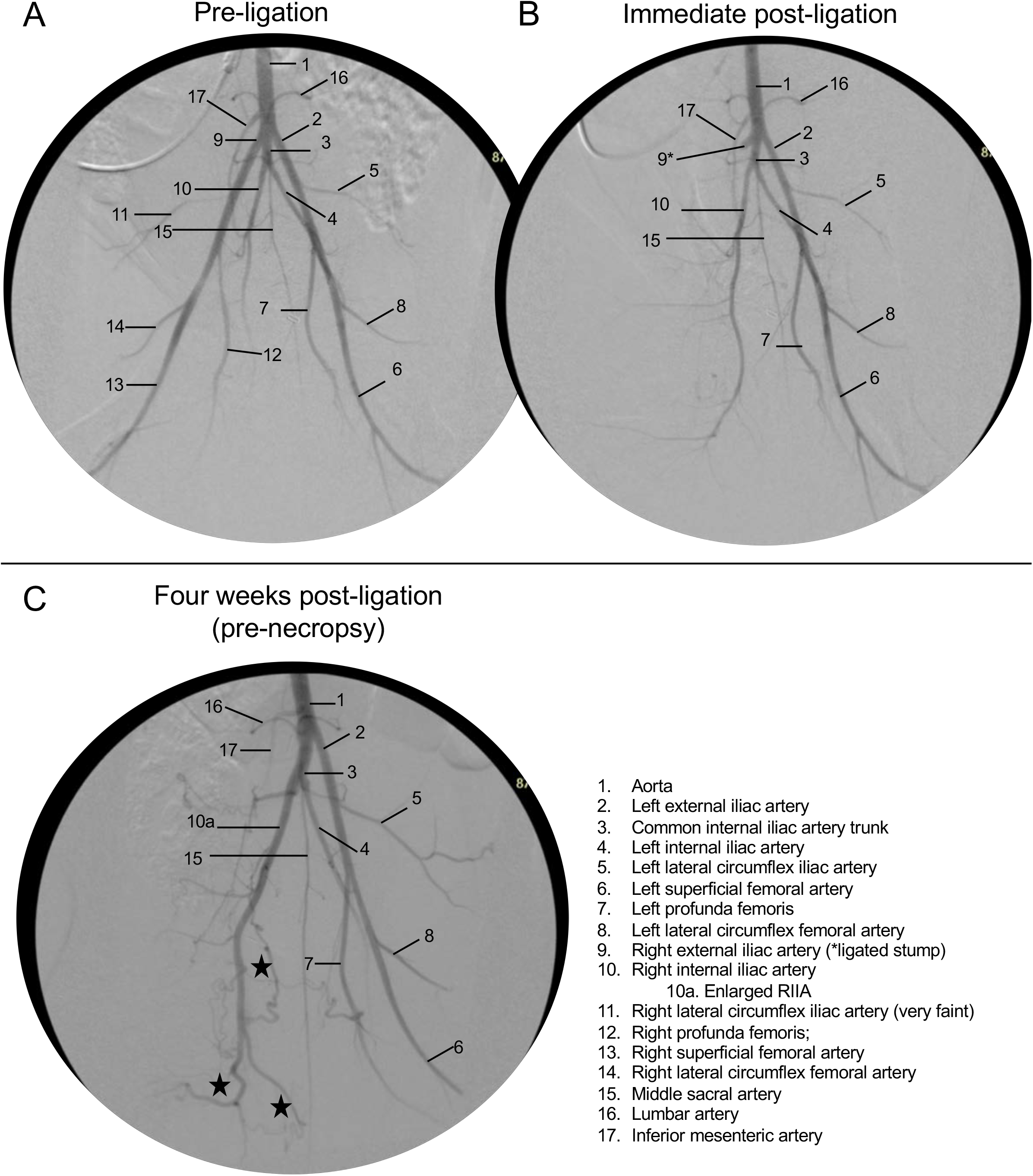
Aortogram with runoff angiography, Version 1. Sample full videos are available in the Supplementary Material. **(A)** Pre-ligation (baseline). **(B)** Immediate post-ligation angiogram. The REIA and all distal vessels are absent. **(C)** Four weeks later, pre-necropsy. Stars indicate collaterals from an enlarged RIIA which reconstitute the SFA and PFA (refer to above videos).

**Table 3.**
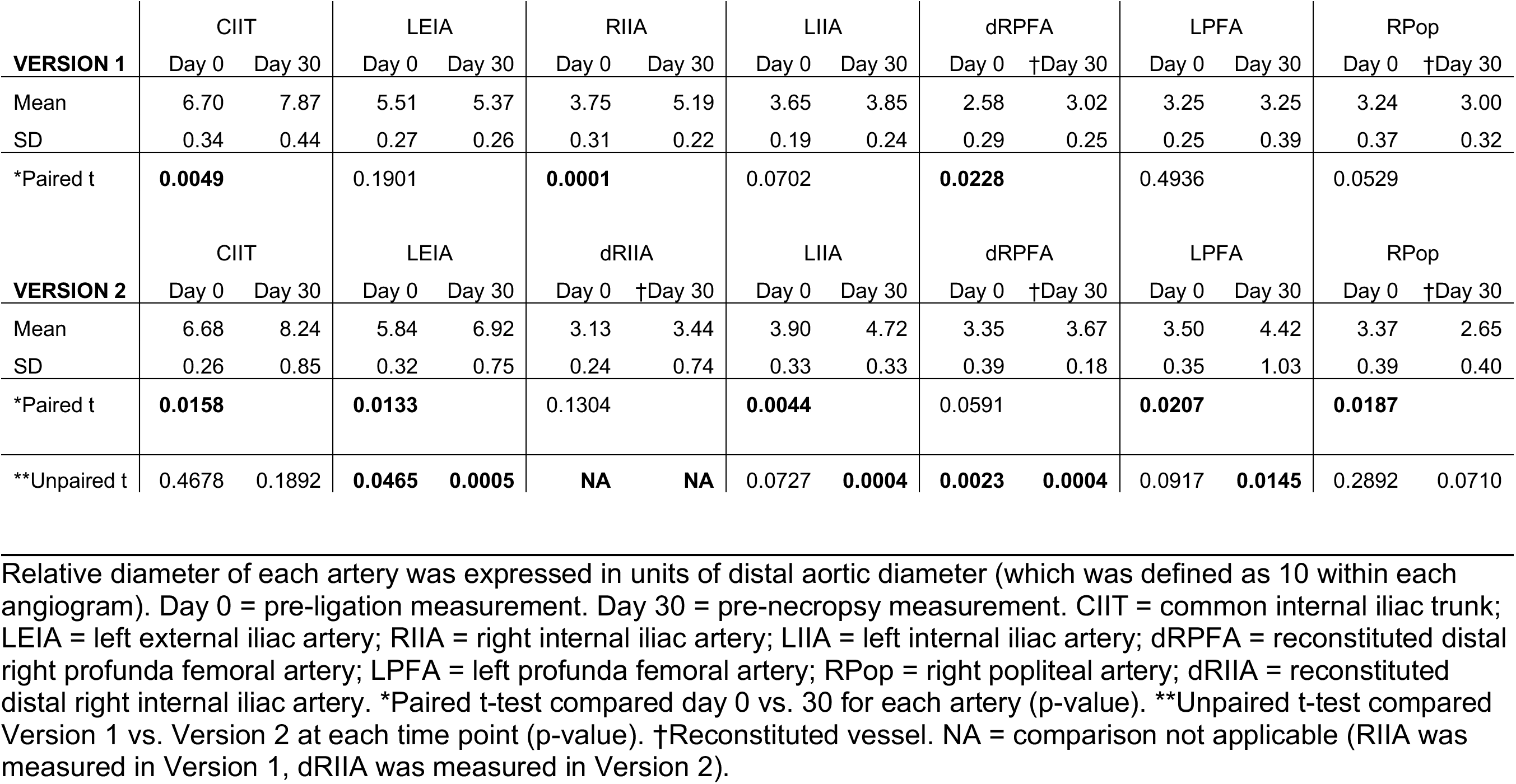
Relative hindlimb arterial diameters, Version 1 vs. 2, at days 0 vs. 30.

**Fig. tf05.**
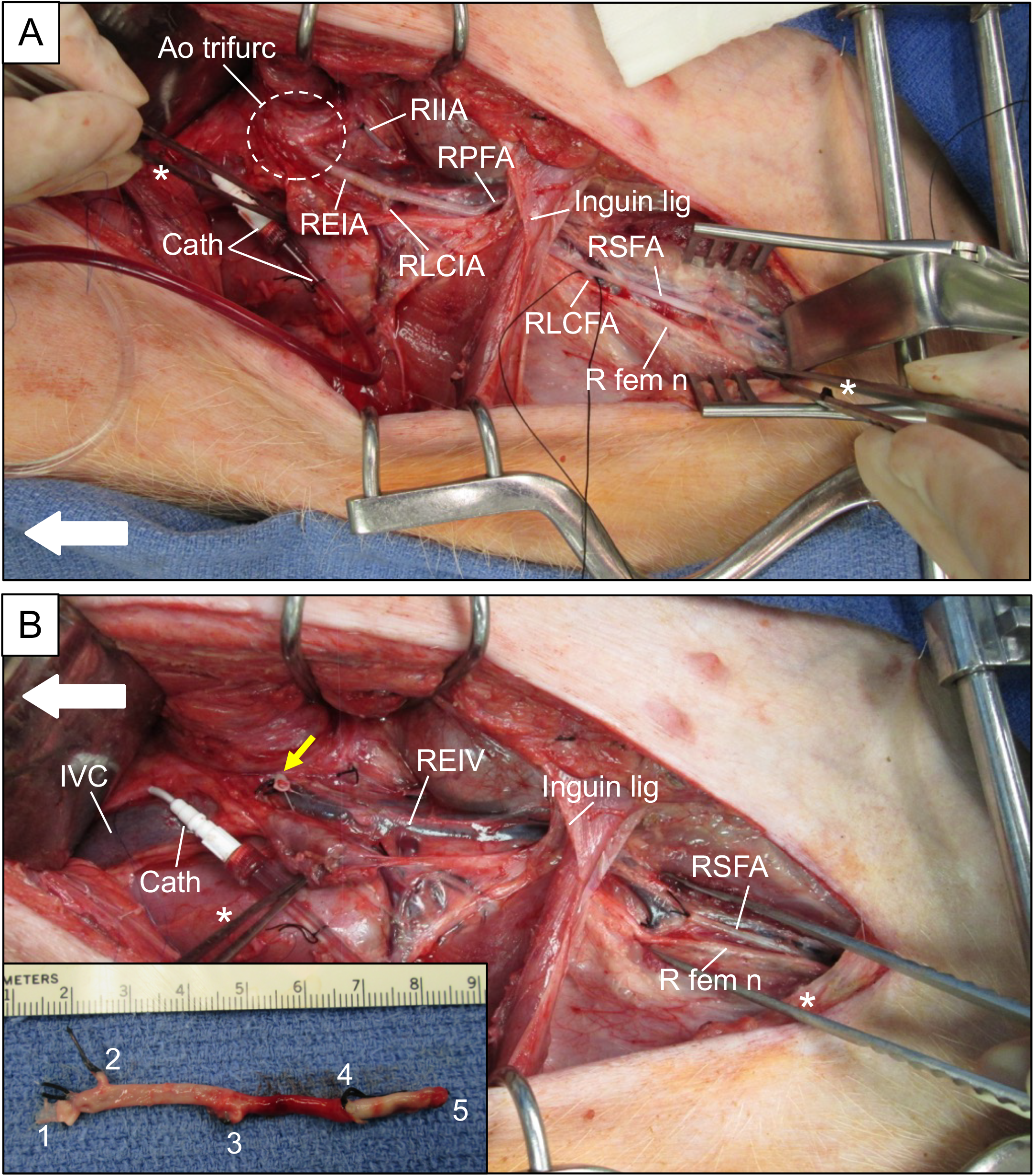
Arterial Interuption procedure (Version 2), intraoperative photos. **(A)** Prior to ligation, showing a right retroperitoneal dissection. Large white arrow points cephalad. The peritoneal sac, containing the intraabdominal viscera, was retracted medially and cephalad. The right iliofemoral complex has been dissected out above and below the inguinal ligament (Inguin lig). The region of the aortic trifurcation (Ao trifurc) is indicated with a dashed white circle. The right internal iliac artery (RIIA) already has been ligated (only for Version 2). There is a silk loop around the right lateral circumflex femoral artery (RLCFA). Cath = angiography cather & line; REIA = right externial iliac artery; RLCIA = right lateral circumflex iliac artery; RPFA = right profunda femoris artery; RSFA = right superficial femoral artery; R fem n = right femoral nerve. **(B)** The right iliofemoral complex has been ligated and excised down to the distal RSFA, with ligation of the RPFA and RCIA. The right external iliac vein (REIV) is visible in the bed of the resected artery. The yellow arrow indicates the stump of the ligated REIA. DeBakey forceps indicated with asterisk (*). **Inset**: resected right iliofemoral complex. Vessel stumps: 1 = REIA; 2 = RLCIA; 3 = RPFA; 4 = RLCFA; 5 = RSFA.

**Fig. tf06.**
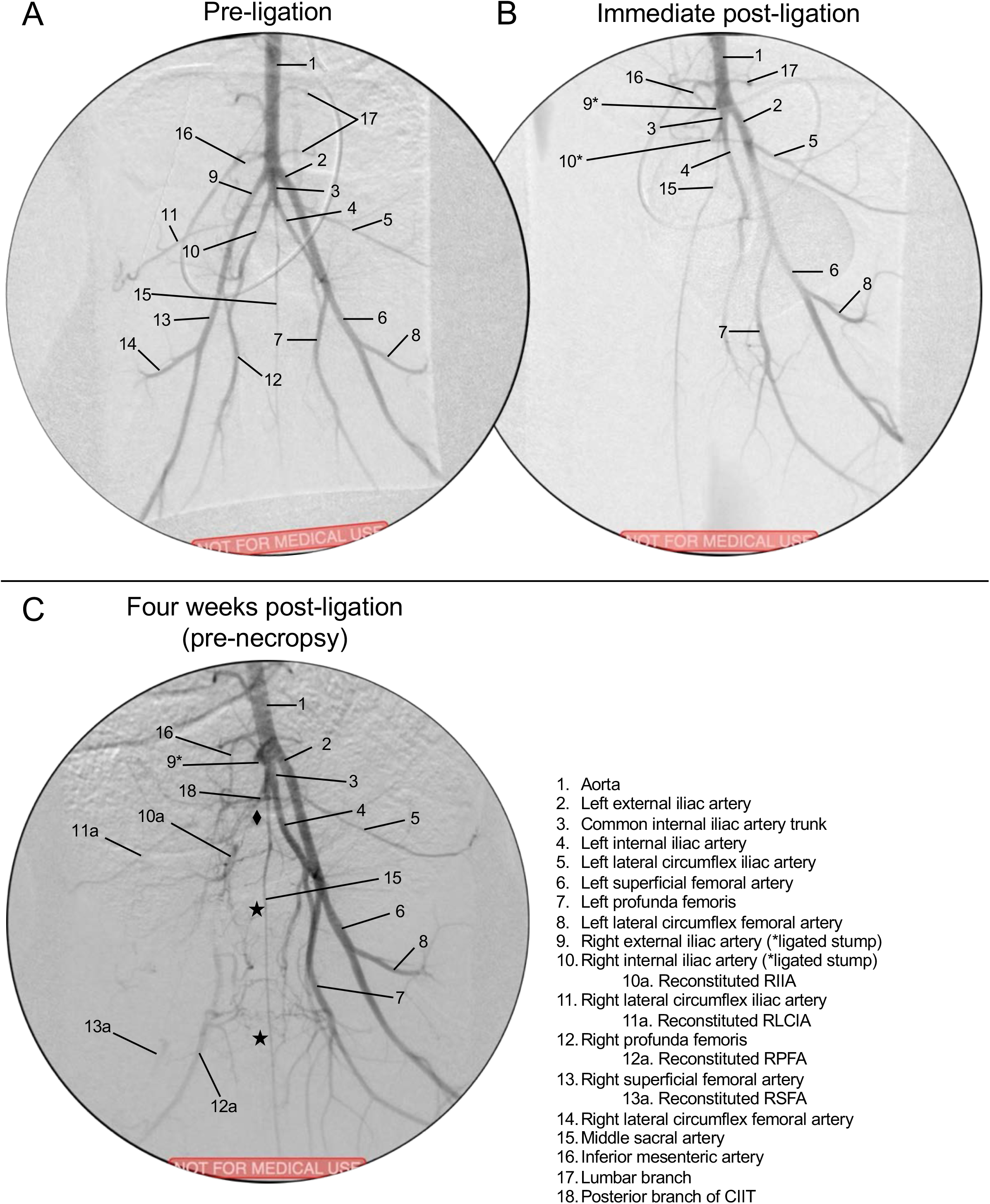
Aortogram with runoff angiography, **Version 2.** Refer to full videos in the Supplementary Material. **(A)** Pre-ligation (baseline). **(B)** Immediate post-ligation angio. The REIA, RIIA, and all distal vessels are absent. **(C)** Four weeks later, pre-necropsy. Stars (★) indicate cross-over collaterals from LPFA which reconstitute the RPFA and then the RSFA; diamond (♦) indicates collaterals from CIIT/ LIIA which reconstitute the RIIA.

### Arteriogenesis

The relative arterial diameters of the hindlimb arteries at pre-ligation (day 0) and pre-necropsy (day 30) are shown in Table 2. Prior to ligation (i.e., day 0), the mean diameter of the LEIA and the dRPFA (distal right profunda femoral artery); were 6% and 30% larger, respectively, in the Version 2 compared to the Version 1 subjects (p< 0.05). The meaning of this difference in baseline arterial diameter between Version 1 vs. 2 subjects is not clear. After 30 days of ischemia in Version 1 subjects, the CIIT, RIIA, and dRPFA increased in diameter by 17%, 38%, and 17%, respectively (p < 0.05); the LIIA and RPop both trended toward increased diameter, but did not reach significance. The dRPFA and RPop both represented reconstituted vessels at 30 days. After 30 days of ischemia in Version 2 subjects, the CIIT, LEIA, LIIA, and LPFA all had a diameter increase in the range of 20% (23%, 18%, 21%, and 26%, respectively; p < 0.05). The dRPFA trended toward increased diameter, but this did not reach significance. The only vessel noted to have a decrease in mean diameter at 30 days was the RPop in the Version 2 subjects, which decreased by 21% (p < 0.05).

### Ischemic Myopathy

Chronically ischemic muscle from both porcine and human subjects exhibited microscopic features of ischemic myopathy (Fig. tf08). H&E micrographs of chronically ischemic (day 30) vs. control (contralateral) porcine gastrocnemius muscle are shown in Fig. tf08A & C, respectively, along with analogous micrographs of gastrocnemius muscle from a claudicating PAD patient and a control non-PAD patient (Fig. tf08B & D, respectively). Control muscle from both swine (Fig. tf08A) and humans (Fig. tf08B) demonstrated polygonal myofibers having relatively similar shape and size, with barely perceptible endomysium and perimysium. However, in ischemic muscle from both swine (Fig. tf08C) and humans (Fig. tf08D), there was a wide range of myofiber size and shape (demonstrating significant myofiber degeneration), with thickening of the endomysium and perimysium (demonstrating significant fibrosis). The myopathic features and other characteristics of chronically ischemic muscle from PAD patients have been previously described [*13–16*].

**Fig tf08.**
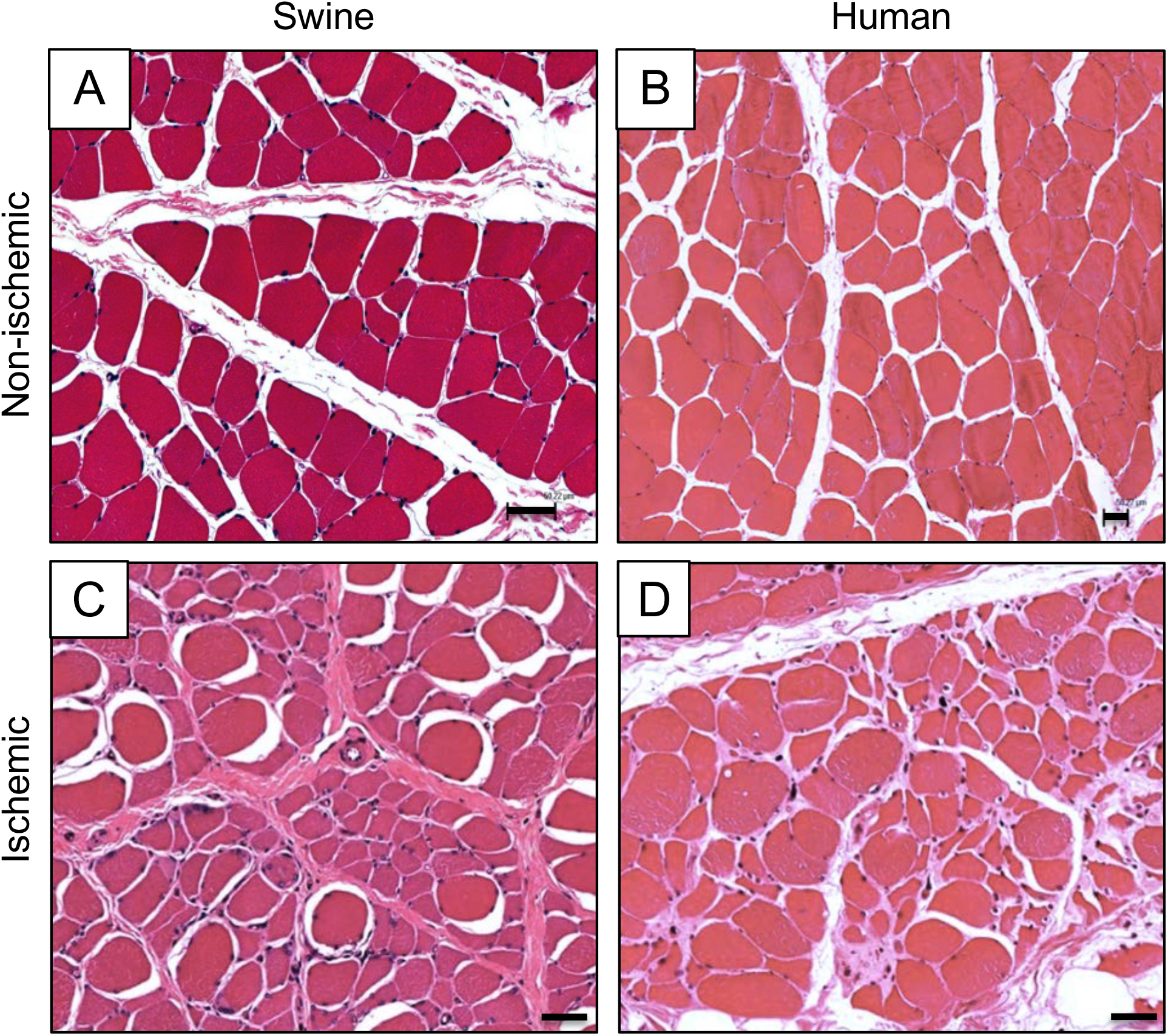
Ischemic myopathy in porcine vs. human subjects. **(A & B)** Non-ischemic muscle, human vs. porcine, respectively. **(C & D)** Ischemic muscle, human vs. porcine, respectively. Samples from both human and porcine subjects were taken from the medial head of the gastrocnemius muscle. H&E; bars = 50 µm.

## DISCUSSION

Herein we have described the pattern and relative size of the collateral pathways that develop after ligation of the native iliofemoral pedicle in a porcine model of PAD. These data demonstrate that two versions of hindlimb ischemia induction (right iliofemoral artery ligation/excision with and without right internal iliac artery ligation) produce differing collateral pathways along with outward remodeling/arteriogenesis (seen as increases of the diameter) of the major inflow vessels. We found that after ligation and excision of the right iliofemoral artery a dominant pathway developed which connected the patent RIIA to the reconstituted RPFA and RSFA/popliteal artery of the ischemic limb. However, after ligation and excision of the right iliofemoral artery with concomitant ligation of the RIIA, two co-dominant collateral pathways developed: (i) the first connected the left profunda artery to the reconstituted RPFA which then supplied the reconstituted RSFA/popliteal arteries: (ii) the second connected the common internal iliac trunk and left internal iliac artery to the reconstituted RIIA, which then helped supply the reconstituted RPFA and RSFA/popliteal arteries.

Of note, three strains of swine have been tested to date as models of human PAD: (i) Ossabaw mini-swine fed a high-fat diet in order to mimic metabolic syndrome [*7*]; (ii) Yorkshire swine fed regular chow [*8*]; and (iii) domestic swine fed regular chow (present report). Our data has confirmed previous findings from the groups of Drs. Brewster and Lefer [*7,8*] showing that all of these models, which have utilized both endovascular and open arterial interruption for induction of hindlimb ischemia, have demonstrated the ability to recapitulate the pathophysiologic characteristics of human PAD, including decreased hindlimb hemodynamics/ perfusion and ischemic myopathy (i.e., end-organ damage). Our study also demonstrates that ligation of the iliofemoral artery, in addition to producing a persistently decreased ABI at rest, also produces a significant decrease in the oxygen saturation of the gastrocnemius that is persistent out to 30 days. The use of domestic swine fed regular chow along with an open arterial ligation technique (present report) allows for a relative cost saving (a less expensive animal and diet, and lower cost of operative and device supplies). In addition, the open ligation approach does not require specialized endovascular experience or operative setup, yet retains the ability to tailor the degree of ischemia. The main limitation of the open ligation approach compared to the one using endovascular occlusion [*7,8*] is the requirement for a large incision for the arterial ligations. The degree of tissue trauma and pain associated with this open procedure might affect the recovery of the subjects from the induction operation, and may become a confounding factor when studying functional status (e.g., gait, treadmill performance) of the subjects.

A unique aspect of this project was the careful characterization of arteriogenesis and collateral formation. We were able to demonstrate specific locations within the porcine vascular network that experienced the greatest luminal growth after induction of ischemia. Quantification of arterial growth at specific sites was used to support the descriptive finding of reproducible collateral formation that were observed in both versions of hindlimb ischemia during angiography. Arteriogenesis is thought to be a reactive process to blood flow redistribution that occurs in occlusive disease [*17–20*]. Changes in fluid dynamics within patent arteries nearby an occlusion cause an increase in shear forces that leads to upregulation of cell adhesion molecules and nitric oxide production in endothelial cells, with a subsequent rise in cytokine and growth factor release, which leads to endothelial and smooth cell proliferation, and a persistent increase (i.e., structural change) in arterial lumen. Given the current understanding of the pathophysiology of the vessel remodeling that occurs in PAD, we can use the swine ischemia model for quantitative testing of arteriogenic therapies, as well as to investigate and improve delivery methods and dosing regimens.

Both ligation versions induced a similar amount of ischemia initially and at day 30, per the resting arterial index and muscle StO_2_ data. That is, addition of internal iliac artery ligation did not produce an increase in ischemia that was detectable at rest with the methods we used. Induction of ischemia in the right hindlimb did not affect the arterial pressure index in the contralateral hindlimb. The muscle StO2 in the contralateral control limb appeared to trend downward immediately after induction of right hindlimb ischemia, with apparent recovery by day 30; to be fair, these changes were modest (nonsignificant, p >0.05) and may not have been real. If the muscle StO2 did decrease in the control limb in the absence of change in large artery pressure, this might suggest a remote effect of limb ischemia on small vessel hemodynamics, e.g., through neuromodulation and/or systemic soluble mediators.

Regarding the two premature mortalities in this study, rectal prolapse is common in growing swine from 8-20 weeks old [*21*]. The prolapse in our study may have been exacerbated by increased intraabdominal pressure from postoperative ileus. The subject which died from intestinal ischemia had a stomach distended from recent feeding; this apparently was a consequence of noncompliance with the prescribed 24 h preoperative fast. This noncompliance (i.e., inappropriate access to feeds) appeared to be an isolated event, as other subjects did not have postoperative intraabdominal issues. The development of hoof ulcers in two of the surviving eleven subjects demonstrated that some subjects may have had critical limb ischemia.

Limitations of this study which could impact the validity of our porcine PAD model include its acute onset of ischemia, which differs from the more progressive and chronic nature of human PAD. Another limitation concerns mimicry of clinical conditions; while PAD patients often are aged and have comorbidities such as hypertension, dyslipidemia, and diabetes, our domestic swine were juvenile, without comorbidities. Regarding hemodynamic and perfusion measurements, all data were collected at rest under anesthesia, so relevant exercise-induced phenomena could have been missed. Regarding quantification of arteriogenesis, only larger inflow arteries were evaluated, so growth in smaller, unnamed collateral arteries may also have been missed. Future work can incorporate comorbidities into the model, and/or acquire measurements during stress maneuvers (e.g., post-occlusive hyperemia or exercise).

The development of collateral vasculature is a key mechanism which compensates for arterial occlusion in PAD, and has the potential to be manipulated therapeutically to treat afflicted patients. In particular, the advent of different angiogenic therapies for humans, including administration of angiogenic cytokines (either as recombinant protein or with gene therapy) and, more recently, investigations of stem/progenitor cell therapy, has opened a new frontier in therapeutic opportunities for PAD [*22–24*]. We believe that the present report can serve as a reference for future studies on novel therapeutic interventions that would enhance collateral growth. For example, a porcine model could help in the optimization of delivery methods, dosing and efficacy testing of experimental arteriogenic treatments, including cell-based therapies [*23,25-28*].

## CONCLUSION

We documented collateral network formation and quantified arteriogenesis in our open model of porcine hindlimb ischemia model. These phenomena are important compensatory mechanisms in the ischemic lower extremity of the patient with symptomatic PAD. Using the porcine model and the measurements described herein, we intend to develop, study, and optimize therapies for the treatment of PAD, particularly for patients whose revascularization options are limited.

## Supporting information

Vid 1

Vid 2

Vid 3

Vid 4

Vid 5

Vid 6

Vid 7

Vid 8

Vid 9

Vid 10

Vid 11

Vid 12

## Abbreviations

AAALAC: Association for Assessment and Accreditation of Laboratory Animal Care International
ABI: Ankle/Brachial Index
CLI: Chronic Limb Ischemia
CIIT: Common Internal Iliac Trunk
DICOM: Digital Imaging and Communications in Medicine
dRPFA: reconstituted distal Right Profunda Femoral Artery
dRIIA: reconstituted distal Right Internal Iliac Artery
IACUC: Institutional Animal Care and Use Committee
LEIA: Left External Iliac Artery
LIIA: Left Internal Iliac Artery
LPFA: Left Profunda Femoral Artery
REIA: Right External Iliac Artery
RLCIA: Right Lateral Circumflex Iliac
RLCFA: Right Lateral Circumflex Femoral Artery
RIIA: Right Internal Iliac Artery
RPFA: Right Profunda Femoral Artery
RSFA: Right Superficial Femoral Artery
RPop: Right Popliteal Artery
PAD: Peripheral Arterial Disease
Moxy: Muscle Oximetry
StO_2_: Muscle oxygen saturation
THb: Total Hemoglobin

## ACKNOWLEDGEMENTS

This work was supported by NIH grants R01 AG034995 and R01 AG049868, by the Charles and Mary Heider Fund for Excellence in Vascular Surgery, by internal seed funds from the Department of Surgery of the University of Nebraska Medical Center, and with resources and the use of facilities at the VA Nebraska-Western Iowa Health Care System. Portions of this study were presented at Vascular Discovery: From Genes to Medicine (San Francisco, CA; May, 2018). The authors would like to acknowledge the technical assistance of Chris Hansen.

## SUPPLEMENTAL MATERIAL

Figures tf09S-tf13S. Videos

1. “Vid01_v1_PreLigAngio_Proximal.mp4”. Description: Version 1 aortobifemoral angiography shot pre-ligation, proximal view.
2. “Vid02_v1_PreLigAngio_Distal.mp4”. Description: Version 1 aortobifemoral angiography shot pre-ligation, distal view.
3. “Vid03_v1_PostLigAngio_Proximal.mp4”. Description: Version 1 aortobifemoral angiography shot immediately post-ligation, proximal view.
4. “Vid04_v1_PostLigAngio_Distal.mp4”. Description: Version 1 aortobifemoral angiography shot immediately post-ligation, distal view.
5. “Vid05_v1_PreNecAngio_Proximal.mp4”. Description: Version 1 aortobifemoral angiography shot immediately pre-necropsy (30 days after ligation), proximal view.
6. “Vid06_v1_PreNecAngio_Distal.mp4”. Description: Version 1 aortobifemoral angiography shot immediately pre-necropsy (30 days after ligation), distal view.
7. “Vid07_v2_PreLigAngio_Proximal.mp4”. Description: Version 2 aortobifemoral angiography shot pre-ligation, proximal view.
8. “Vid08_v2_PreLigAngio_Distal.mp4”. Description: Version 2 aortobifemoral angiography shot pre-ligation, distal view.
9. “Vid09_v2_PostLigAngio_Proximal.mp4”. Description: Version 2 aortobifemoral angiography shot immediately post-ligation, proximal view.
10. “Vid10_v2_PostLigAngio_Distal.mp4”. Description: Version 2 aortobifemoral angiography shot immediately post-ligation, distal view.
11. “Vid11_v2_PreNecAngio_Proximal.mp4”. Description: Version 2 aortobifemoral angiography shot immediately pre-necropsy (30 days after ligation), proximal view.
12. “Vid12_v2_PreNecAngio_Distal.mp4”. Description: Version 2 aortobifemoral angiography shot immediately pre-necropsy (30 days after ligation), distal view.

**Fig tf09S.**
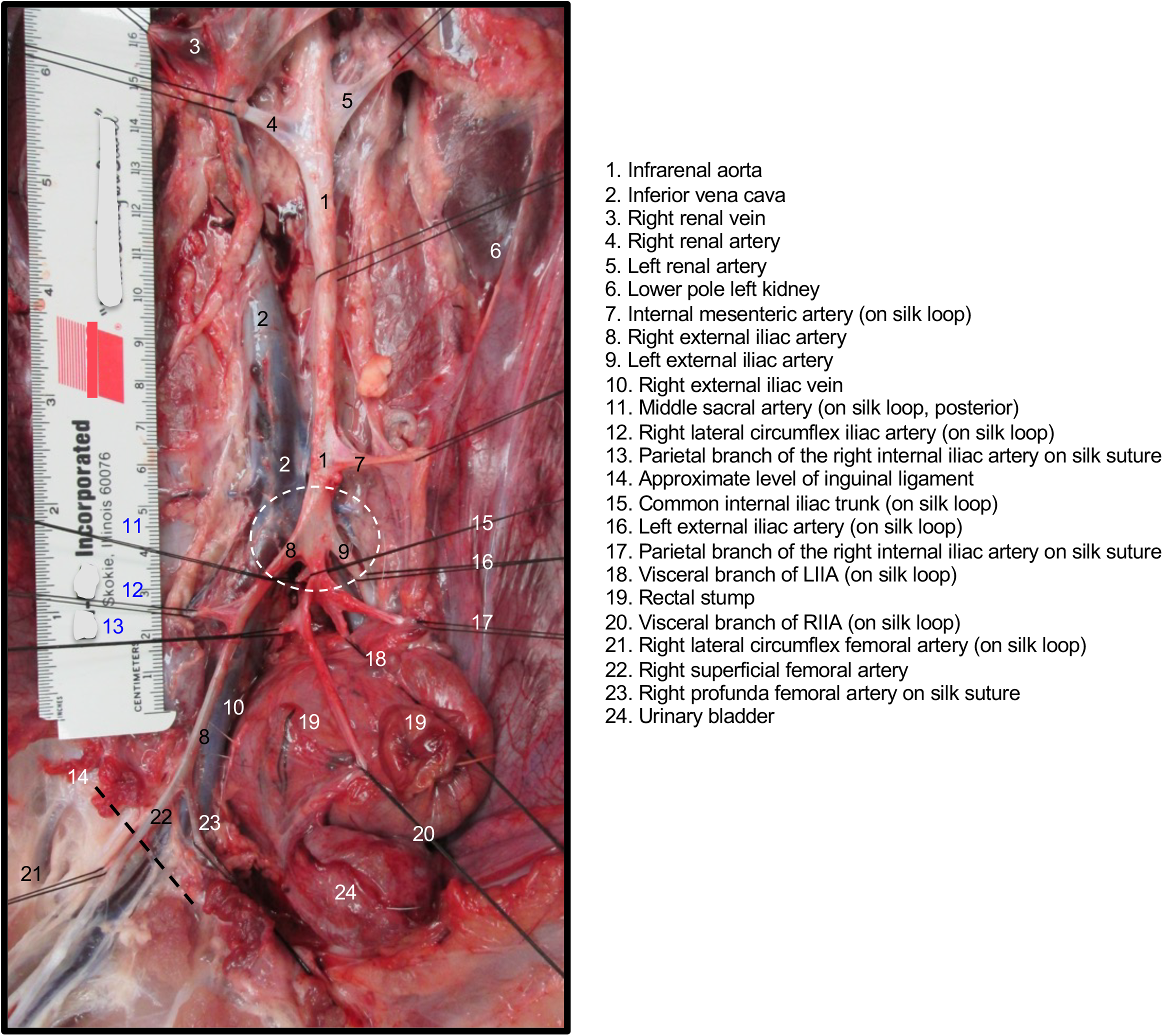
Relevant anatomy of the porcine hindlimb. Nonfixed postmortem dissection of the distal aorta, pelvis, and proximal right hindlimb in the pig. Dashed circle indicates aortic trifurcation.

**Fig. tf10S.**
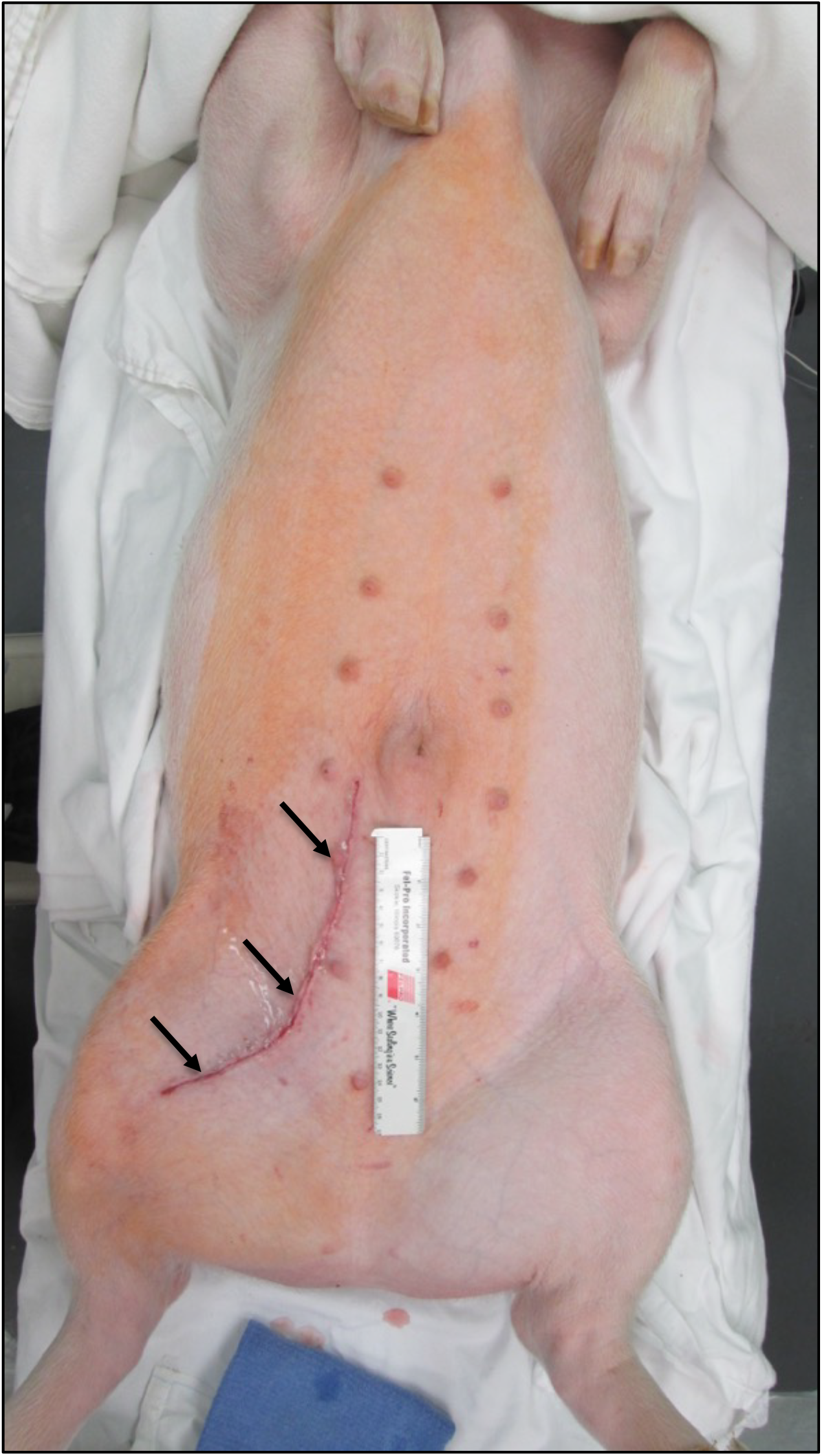
Incision for retroperitoneal dissection. Anesthetized porcine subject is supine, with overhead view of ventral chest, abdomen, and pelvis; cephalad is toward top of image. Freshly closed incision indicated with arrows. Ruler is 15 cm long.

**Fig tf11S.**
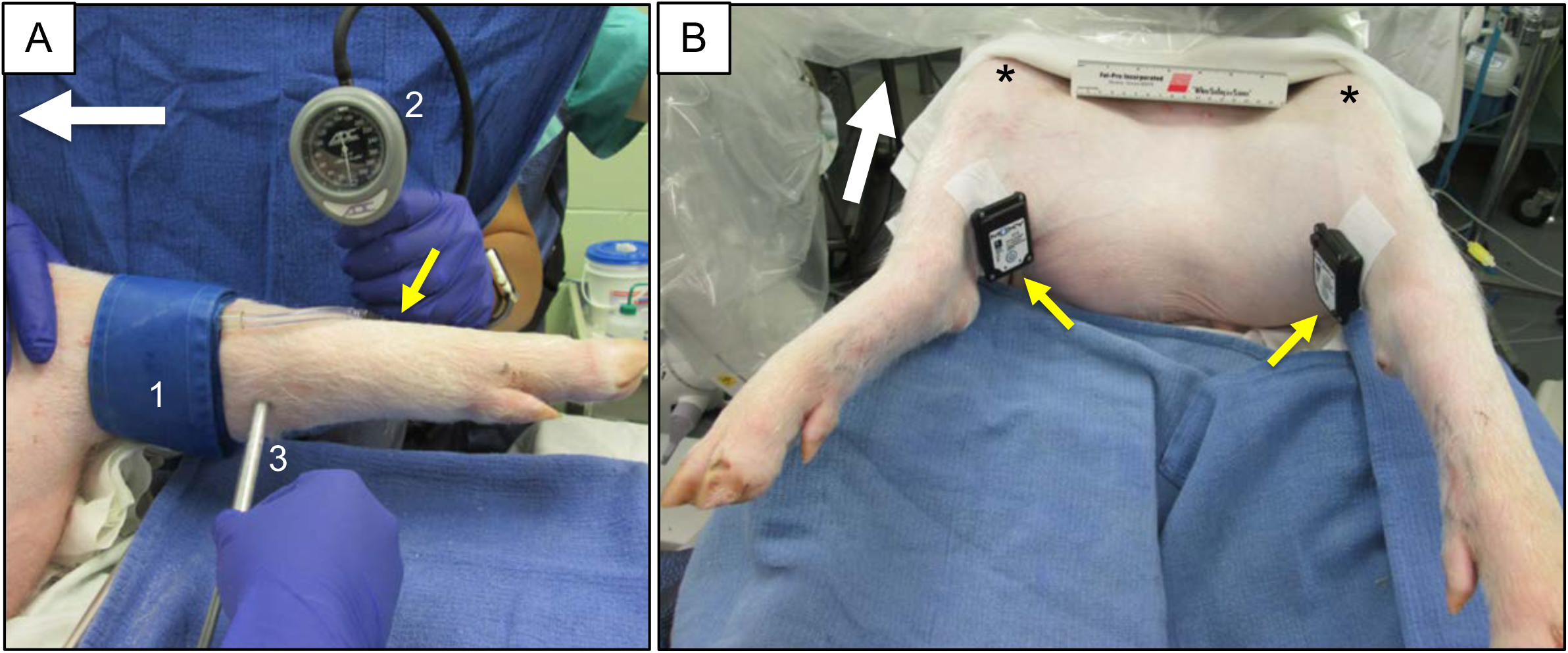
Measurements of ABI and Muscle oxygen saturation in pigs. **(A)** A pediatric blood pressure cuff (1) has been positioned just proximal to the calcaneous of the left hindlimb for determination of arterial pressure (2). The doppler probe (3) has been placed over the posterior tibialis artery. Alternatively, the doppler probe can detect arterial signal over the dorsalis pedis artery (small yellow arrow). Large white arrow = pointing cephalad. **(B)**. Transcutaneous measurement of muscle hemoglobin/myoglobin oxygen saturation (Moxy). The wireless Moxy instrument probes (small yellow arrows) have been placed over each medial calf (medial gastrocnemius muscle). Prior to measurement acquisition, the Moxy probes were covered with a loose dark wrap (black 3M™ Coban™) to minimize interference from ambient room light. Inferior view of the supine porcine subject (a castrated male); each stifle (knee) joint indicated with an asterisk (*); large white arrow = pointing cephalad.

**Fig tf14S.**
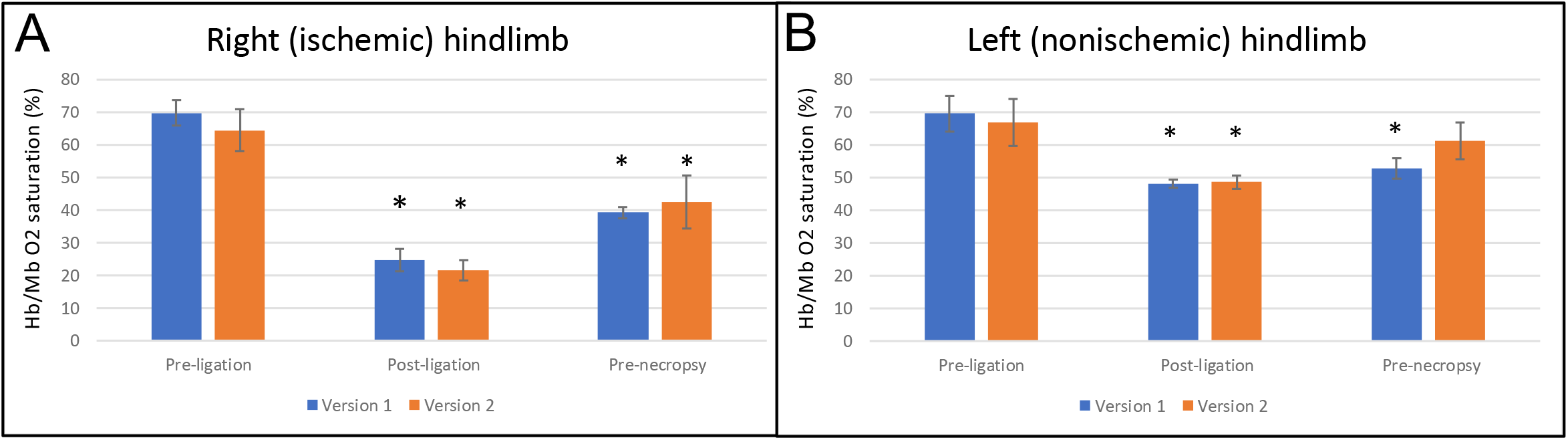
Muscle hemoglobin/myoglobin oxygen saturation (StO_2_ or Moxy), raw readings. Right ischemic hindlimb (Panel A) and left no-ischemic hindlimb (Panel B). Each bar represents mean ± sd of all measurements (raw saturation value) from each subject at each time point; measurements were taken at a constant location (medial gastrocnemius) at each time point. *p < 0.05 compared to pre-ligation.

